# Adipocyte sphingosine kinase 1 regulates histone modifiers to disrupt circadian function

**DOI:** 10.1101/2024.09.13.612486

**Authors:** Andrea Anderson, Anna Kovilakath, Maryam Jamil, Johana Lambert, L. Ashley Cowart

## Abstract

Circadian rhythms align biological functions with the 24-hour day-night cycle, but modern artificial light disrupts these patterns, contributing to health issues like obesity and cardiovascular disease. The circadian clock operates through a transcriptional-translational feedback loop involving core components such as BMAL1 and CLOCK. Recent research has shown circadian variations in sphingolipid metabolism, specifically sphingosine-1-phosphate (S1P), which plays crucial signaling roles. This study investigates the sphingolipid enzyme, sphingosine kinase 1 (SphK1), which converts sphingosine to S1P, as a circadian-regulated gene in adipocytes. We find that SphK1 expression and activity follow a circadian rhythm, regulated by BMAL1 and CLOCK binding to its promoter. Adipocyte-specific SphK1 knockout mice exhibit disrupted circadian rhythms, and impaired adipocyte function. Additionally, SphK1 deficiency leads to reduced histone acetylation and altered histone deacetylase (HDAC) localization, affecting gene regulation. These results highlight the critical role of SphK1 in linking lipid metabolism with circadian biology.

## Introduction

Circadian rhythm research focuses on the study of biological processes that align with the 24-hour day, having evolved to anticipate and adapt to daily cycles of light and dark caused by Earth’s rotation as it orbits around the sun (1, 2). Humans experience a highly disruptive circadian pattern in today’s 24/7 artificially-lit society, leading to multiple pathological manifestations (3, 4). At the molecular level, the circadian clock is governed by a transcriptional-translational feedback loop for the duration of approximately 24 hours. The core mechanism involves heterodimerization of transcription factors, BMAL1 and CLOCK, which in turn bind E-box elements to activate transcription of PERIODs (PER1-3) and CRYPTOCHROME (CRY1-2) in the negative core loop; REV-ERBs (α, β), in the negative accessory loop; and RORs (α, β, γ), in the positive accessory loop. In typical negative feedback loop mechanism, as PER and CRY accumulate, they heterodimerize, translocate into the nucleus and repress BMAL1:CLOCK activity. This in turn decreases PER, and CRY due to lack of transcription as well as coordinated ubiquitination-mediated degradation. Once PER:CRY complexes are degraded, a new cycle of BMAL1:CLOCK-mediated transcription begins (5).

Findings in the last decade point to circadian oscillations in the sphingolipid pathway (6-8). Sphingolipids are a class of bioactive lipids involved in cell survival, apoptosis, growth, proliferation, and differentiation (9). *De novo* biosynthesis of sphingolipids occurs when serine palmitoytransferase catalyzes a free fatty acid and amino acid (10). Ultimately, a delicate balance is achieved among the levels of these lipid structural and signaling metabolites in the cell, including: sphingomyelin, ceramide, glycosphingolipids, ceramide-1-phosphate, sphingosine, and sphingosine-1-phosphate (S1P).

This study focuses on sphingosine kinase 1 (SphK1), an enzyme that phosphorylates sphingosine to generate the bioactive signaling sphingolipid, S1P, capable of a wide variety of functions, ranging from intracellular to autocrine and paracrine signaling (9). Our lab previously identified a role for adipocyte SphK1 in the maintenance of liver homeostasis, highlighting the impact of adipose-liver crosstalk (11). Furthermore, it is well established that histone modifications and their circadian regulation are highly coordinated in the liver to affect numerous metabolic outcomes, such as cholesterol synthesis and secretion (12-14). Prior research also suggests that S1P can modulate the activity of histone-modifying enzymes (15, 16).

Based on these findings, we hypothesized that SphK1 is a circadian-regulated gene in adipocytes and that loss of SphK1 may disrupt adipocyte circadian functions. Our study confirms that SphK1 expression follows a circadian rhythm, suggesting it plays a vital role in maintaining the metabolic and physiological balance within adipocytes. Sphk1 expression exhibits circadian oscillations in mouse fibroblast cells, lung tissue, and adipocytes, driven by the binding of circadian proteins BMAL1 and CLOCK to specific E-box elements in its promoter. The enzymatic activity of SphK1 also follows a circadian rhythm, indicating a time-dependent regulation of its function. We have shown that mice lacking SphK1 in adipocytes (SK1^fatKO^) results in impaired circadian rhythms, and abnormal adipocyte function. Furthermore, SK1^fatKO^ adipocytes exhibit reduced histone acetylation, impairing transcriptional activation at circadian gene promoters, though S1P supplementation can partially restore these epigenetic marks. The study also finds that SphK1 deficiency alters histone deacetylase (HDAC) localization and function, further affecting gene regulation. Overall, these findings highlight the importance of SphK1 in maintaining normal circadian clock functions and gene expression in adipocytes, linking lipid metabolism with circadian biology.

## Results

Recent studies have highlighted the circadian regulation of sphingolipid-associated genes. Among approximately 15,000 genes in humans, *Sphk1* was found to oscillate in four human tissues: coronary artery, visceral fat, heart atria and liver (Figure 1A). Similarly, in about 35,000 circadian genetic hits in mice, *Sphk1* was showed oscillatory expression in two systems: the 3T3 fibroblast cell line and lung tissue (Figure 1B). These findings suggest that SphK1 is atleast partially regulated by the circadian clock in humans.

**Figure 1.**
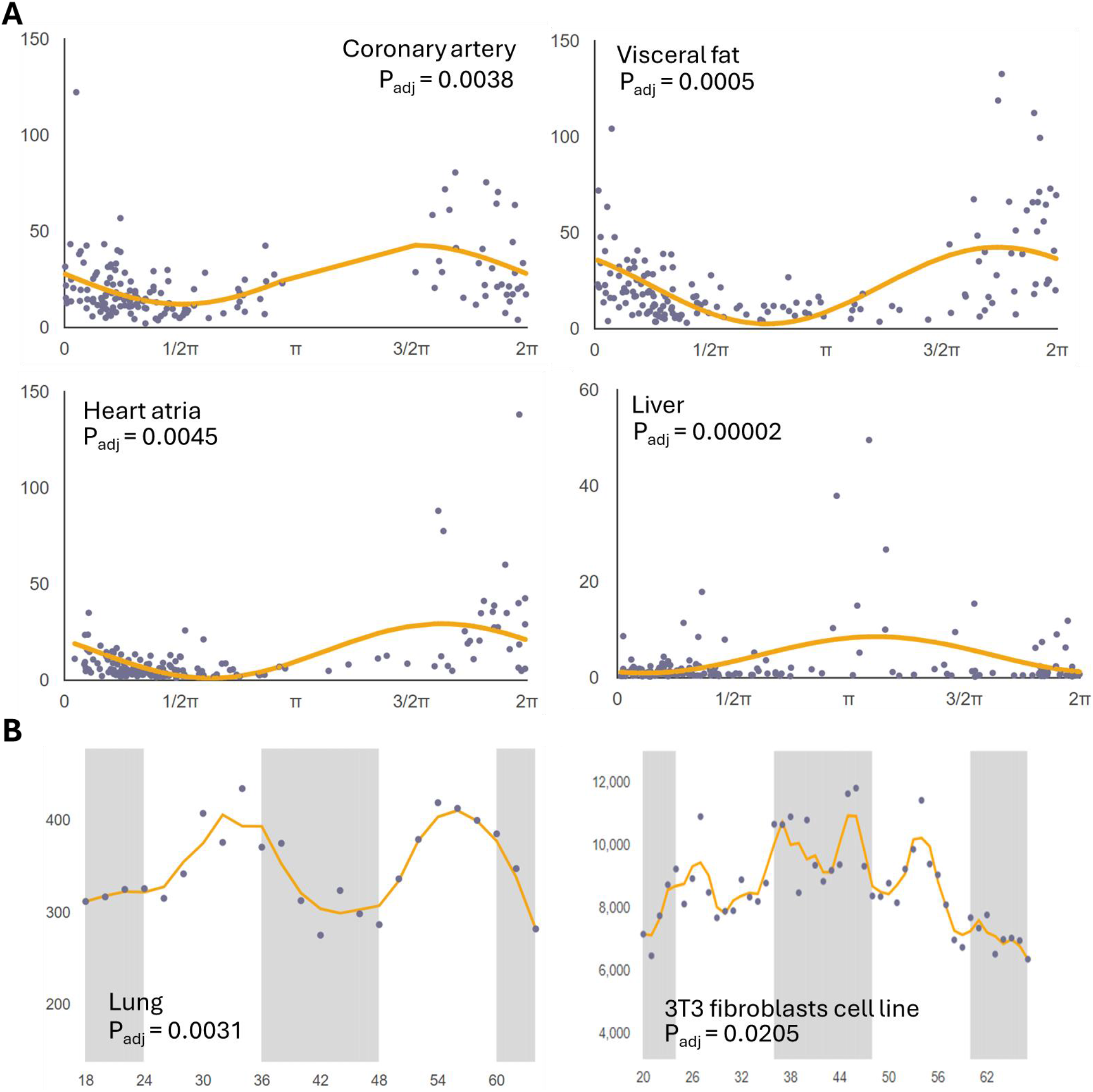
*Sphk1* displays circadian oscillation in human and mouse. *Sphk1* is a fluctuating gene in **A**. human tissue – coronary artery, visceral fat, heart atria, and liver; and **B**. mouse tissue – lung and 3T3 fibroblast cells. Data accessed at http://circadb.hogeneschlab.org.

Given the tight coupling between metabolism and the circadian clock, we hypothesized that SphK1 might be regulated by clock components. A genomic database search for the canonical 5’-CACGTG-3’ E-box element within the SphK1 promoter region identified one E-box approximately 2 kilobases upstream of the transcriptional start site (TSS) and another located about 300 base pairs upstream, proximal to TSS (Figure 2A). In mouse gonadal white adipose tissue (gWAT), *Sphk1* mRNA had a non-constant expression pattern (Figure 2B). In primary adipocytes, *Sphk1* oscillated over a 24-hour period, with a cosinor R^2^ value of 0.76, indicating a robust circadian rhythm. As a control, the canonical negative clock gene *Reverbα* (*Nr1d1*) was also measured and confirmed the validity of the circadian cell culture assay (Figure 2C).

**Figure 2.**
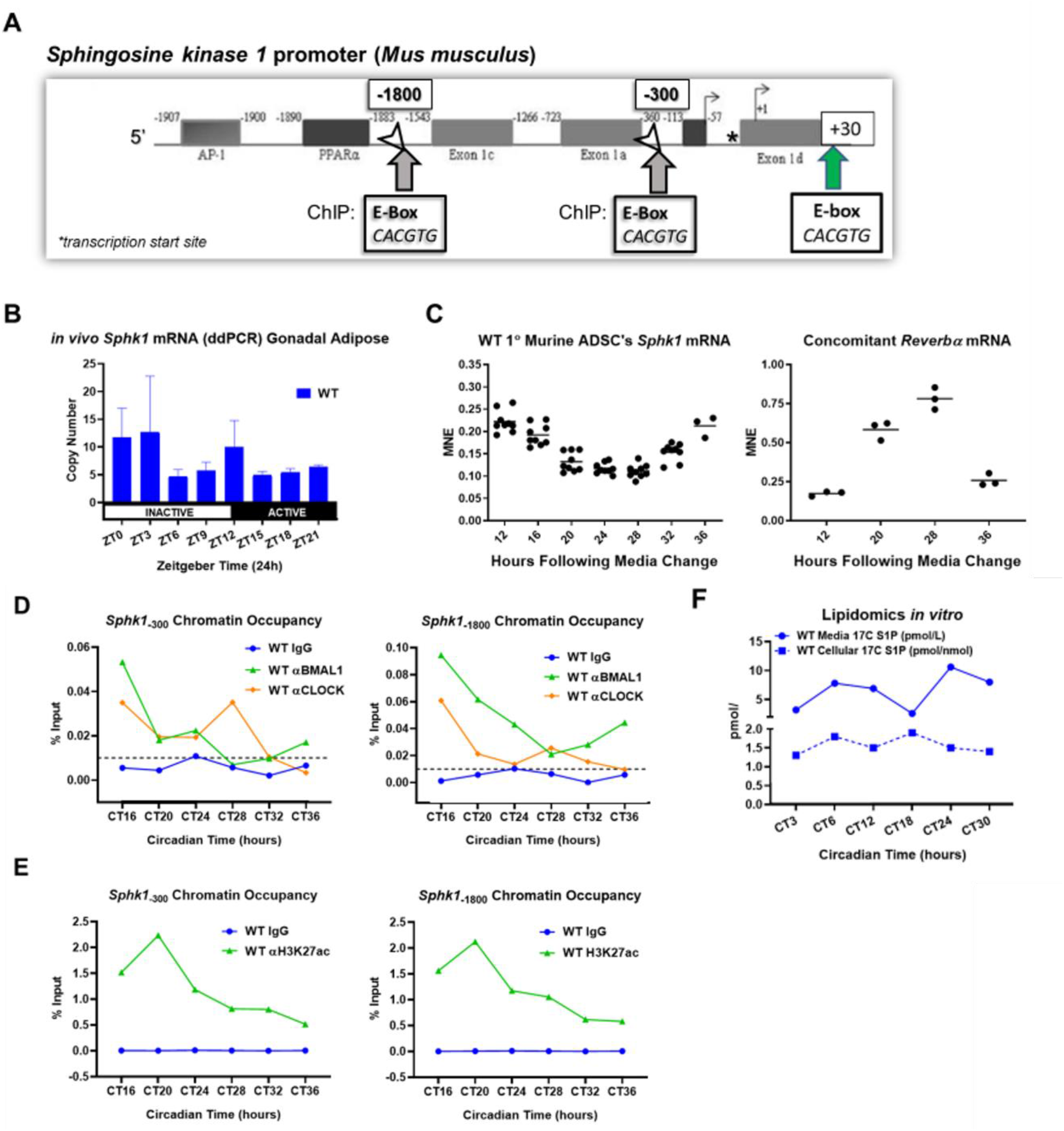
Sphk1 is a circadian gene in adipocytes. **A**. Pictorial representation of the Sphk1 promoter, in which multiple E-boxes reside, including positions -1800 and -300 base pairs from the transcription start site (TSS) (Adapted from Ross and Cowart, et al., 2013). **B**. Sphk1 mRNA expression in gWAT over time in 12hr-light:12hr-dark-entrained wildtype mice. **C**. Sphk1 mRNA expression in primary adipocytes with concomitant expression of Reverbα over 24 hours. **D**. Presence of BMAL1 and CLOCK at the promoter E-box loci in Sphk1 DNA. **E**. H3K27ac ChIP at the Sphk1 promoter E-boxes reveals fluctuations in the occupancy by transcriptionally active chromatin. **F**. Following synchronization, wildtype adipocytes ^17^C-S1P production as measured by LC/MS/MS over a circadian time course when fed 5 µM ^17^C-sphingosine for exactly 30 minutes prior to each harvest. CT, circadian time (CT0 is the time at which synchronization serum shock media is administered); ZT, Zeitgeber time

We next performed chromatin immunoprecipitation (ChIP) analysis to assess the presence of the BMAL1 and CLOCK transactivator complex at the same SphK1 promoter regions. Concomitant pulldowns for acetylated histone 3 lysine 27 (H3K27Ac) were performed as positive controls for transcriptional activation marks. For cell synchronization, growth-arrested mature primary adipocytes were subjected to a serum-shock (50% horse serum and 50% DMEM) for 2 hours, designated as circadian time 0 (CT0) followed by incubation for 46 hours in serum-depleted media. Over a 24-hour time course, ChIP analysis revealed a rhythmic binding of BMAL1, CLOCK, and H3K27Ac to the SphK1 promoter region (Figure 2D). At certain times, signals for core clock transactivator were indistinguishable from background, suggesting BMAL1 and CLOCK regulate *Sphk1* expression, through time-dependent occupancy of the SphK1 E-boxes. To further investigate this phenomenon, lipidomics flux assays to determine whether SphK1 exhibits time-of-day-dependent enzymatic activity were utilized. Cells were treated with a bolus of ^17^C-sphingosine substrate, and turnover to the ^17^C-sphingosine-1-phosphate (^17^C-S1P) product was measured. Consistent with the pulsatile expression of *Sphk1*, the generation of ^17^C-S1P followed a rhythmic pattern (Figure 1E). This suggests SphK1 has time-dependent enzymatic efficacy, or time-dependent enzyme accumulation to handle increased substrate turnover.

The next objective was to characterize the circadian clock in adipocytes lacking SphK1 (SK1^fatKO^) compared to control. A previous study from our lab demonstrated the SK1^fatKO^ mice develop a non-alcoholic fatty liver disease phenotype, and hypertrophic adipocytes with impaired basal lipolysis as a metabolic phenotype (11). Additionally, mRNA sequencing findings of gWAT from control diet-fed WT and SK1^fatKO^ mice revealed that expression of several circadian transcripts were significantly altered (Figure 3).

**Figure 3.**
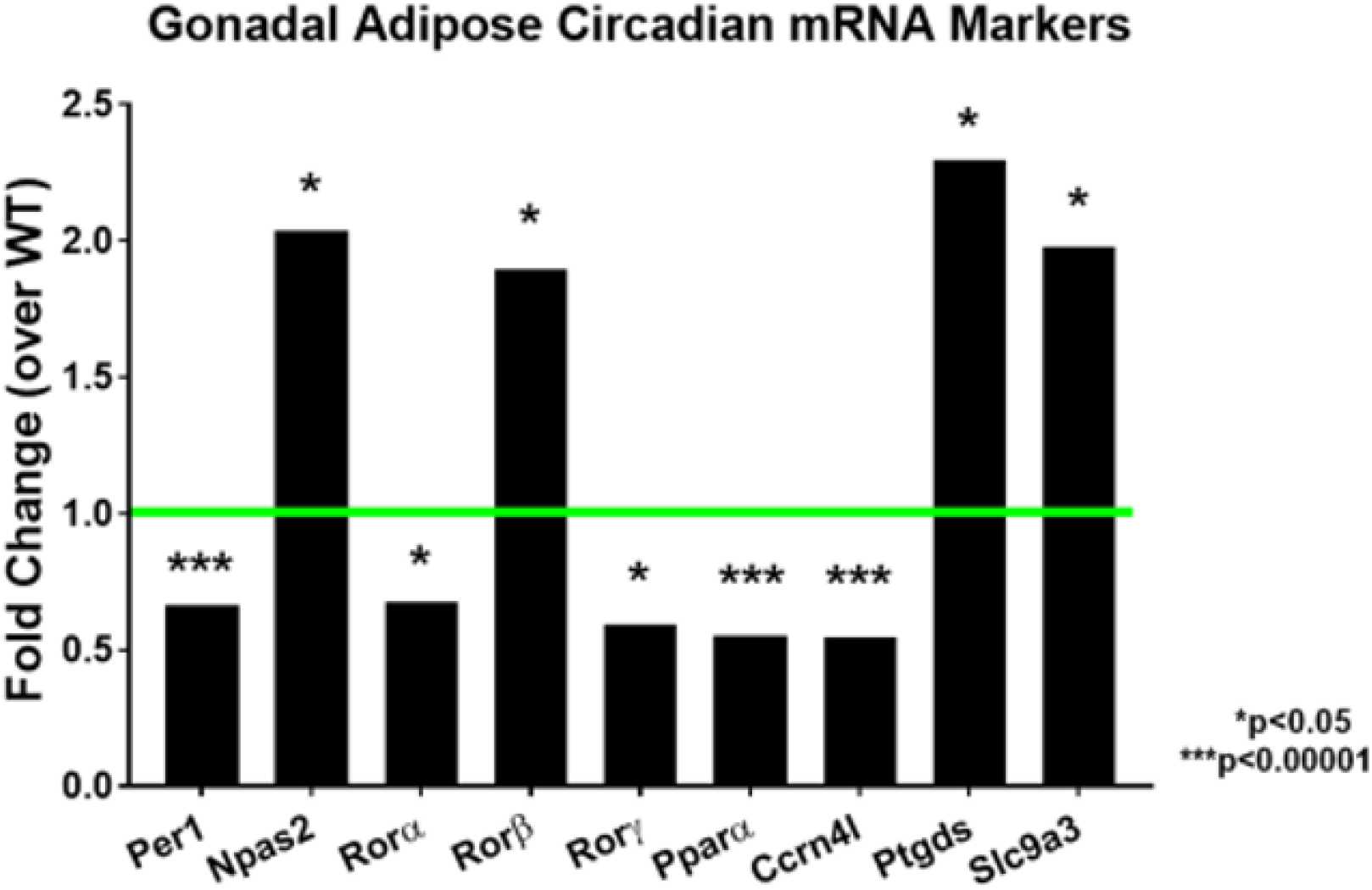
SK1^fatKO^ gWAT mRNA Sequencing analysis. gWAT from pooled samples of male and female C57/bl6J mice (WT or SK1^fatKO^) harvested at ZT9 was analyzed by paired-end fragments per kilobase million mRNA Sequencing and expressed as fold change over wildtype. Ccrn4l, carbon catabolite repression 4-like protein (also known as nocturnin); Npas2, neuronal PAS containing 2; Per1, period 1; Ptgds, prostaglandin D2 synthase; Pparα, peroxisome proliferator activated receptor alpha; Rorα, β, γ, retinoic acid related orphan receptor alpha, beta, and gamma; Slc9a3, solute carrier family 9 member A3. All animals for the dataset were sacrificed at Zeitgeiber Time (ZT) 9 (meaning 9 hours after the light stimulus turns on in the animal housing facility, where 6 a.m. is “lights-on” and ZT0; *i*.*e*., 6 a.m + 9 hours = ZT9) and fasted for 6 hours prior to sacrifice

Based on the mRNA-Seq and *in silico* findings, the next step was to utilize a controlled *in vitro* system for comparing the core clock transcripts in WT and SK1^fatKO^ adipocytes. After synchronization with a horse serum shock, it was observed that certain negative regulators, the *Per* genes displayed a higher amplitude in mutant cells compared to controls (Figure 4A).

**Figure 4.**
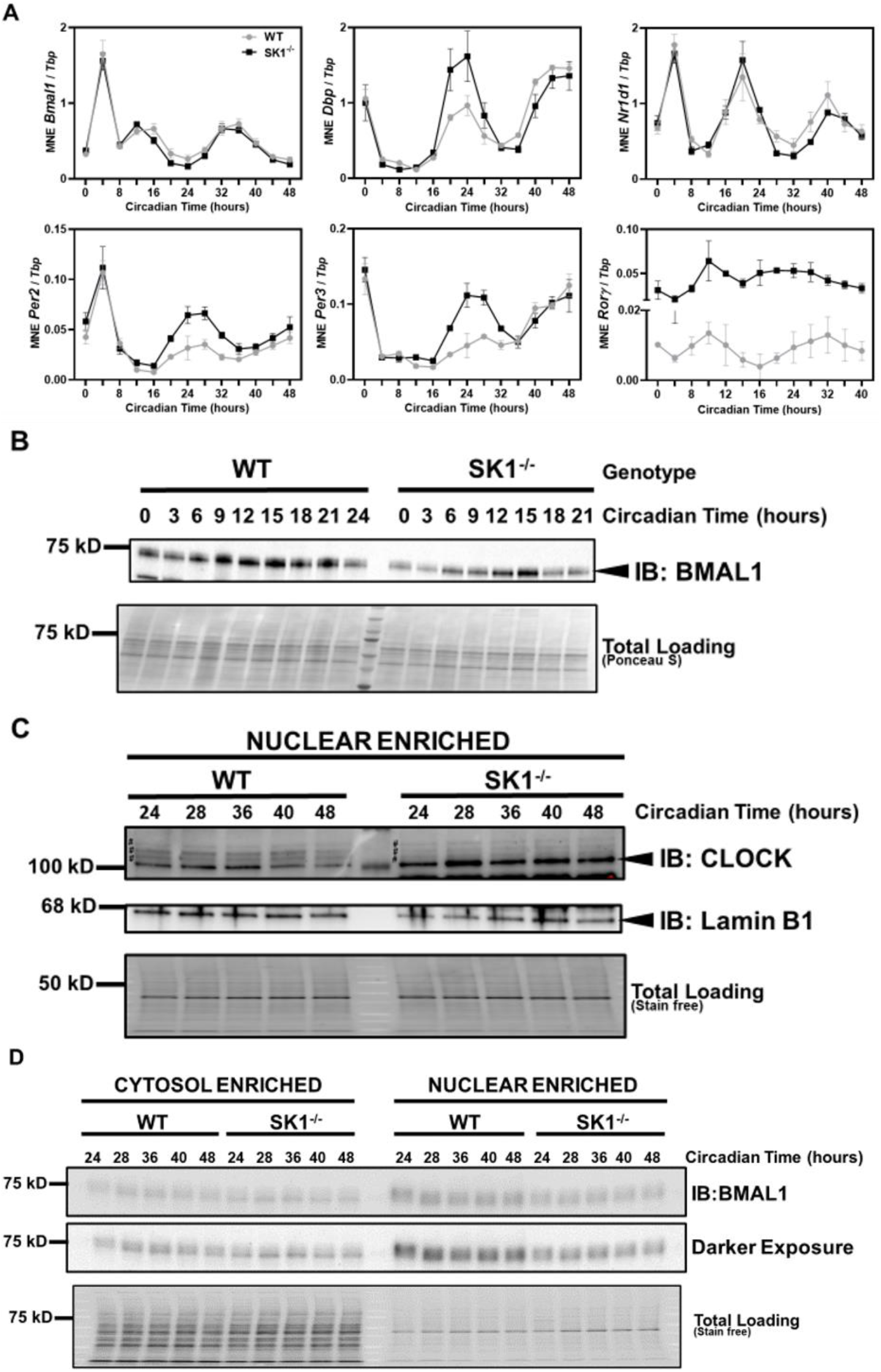
SphK1^-/-^ adipocytes have a dysfunctional circadian clock. **A**. mRNA rhythms of Bmal1 (positive core loop gene), Dbp (canonical clock-controlled gene (ccg)), Nr1d1 (also known as Rev-erbα; negative accessory loop gene), Per2 (negative core loop gene), Per3, and Rorγ (positive accessory loop gene) in synchronized primary matured WT and SPHK1^-/-^ adipocytes over a 48-hour time course. Refer to Figure 2 on page 13 for the molecular circadian clock. **B**. Western blot of BMAL1 protein expression in primary WT and SPHK1^-/-^ matured adipocytes whole cell lysates across the circadian time course. **C**. Expression of BMAL1 protein (by Western blot) in nuclear- and cytoplasmic-enriched fractions of synchronized primary WT and SPHK1^-/-^ matured adipocytes through a time course of 24 hours. **D**. Nuclei of primary WT and SPHK1^-/-^ matured adipocytes were isolated, and CLOCK protein expression over 24 hours was measured by Western blot. # denotes posttranslational modifications. Experiments were performed 3 times in 3 separate preparations of primary cells for culture, and representative blots are shown for the trends that were observed. Bmal1, brain and muscle ARNT-like protein 1; Dbp, D-site of albumin promoter binding protein; Nr1d1, nuclear receptor subfamily 1 group D member 1; Per2, 3, period 2, 3; Rorγ, retinoic acid receptor-related orphan receptor gamma; SK1^-/-^, sphingosine kinase 1 knockout (for brevity in figures).

Moreover, *Dbp* (D-site of albumin promoter binding protein), also demonstrated a higher amplitude in mutant cells compared to control (Figure 4A). Interestingly, the *Bmal1* mRNA rhythm remained unchanged while one of its positive regulators, *Rorγ*, maintained higher gene expression over 48 hours compared to the mutant (Figure 4A). Despite unchanged *Bmal1* mRNA levels, BMAL1 protein expression was lower in SK1^fatKO^ adipocytes, although it still oscillated (Figure 4B). We attribute the spike in *Bmal1* within the first 4 hours to the serum shock synchronization (Figure 4A).

Since BMAL1 is a nuclear translocator, we determined whether cellular localization was affected in SK1^fatKO^ adipocytes. Intriguingly, nuclear expression of BMAL1 protein was diminished compared to controls, suggesting an aberrancy in the clock transactivator machinery (Figure 4C). Conversely, nuclear CLOCK protein levels increased in SK1^fatKO^ adipocytes (Figure 4D). These findings led to the conclusion that SphK1 is necessary for normal adipocyte circadian clock function.

Having established that SphK1 is circadian-regulated and that loss of this gene impairs circadian functionality, the next question was to asess the impact on circadian clock-mediated transcriptional regulation. To explore this, we examined the expression patterns of BMAL1:CLOCK targets by immunoprecipitating components of the transactivator complex at the chromatin E-box regions of *Per1, Reverbα, Dbp*, and *Pparγ* (Figure 5A). In synchronized, non-transfected adipocytes, BMAL1 and CLOCK were detected at circadian time 20 (CT20) at all tested promoters, but CLOCK occupancy at the *Reverbα* promoter was significantly lower in the mutants compared to controls, with similar downward trends observed for the other targets (Figure 5A).

**Figure 5.**
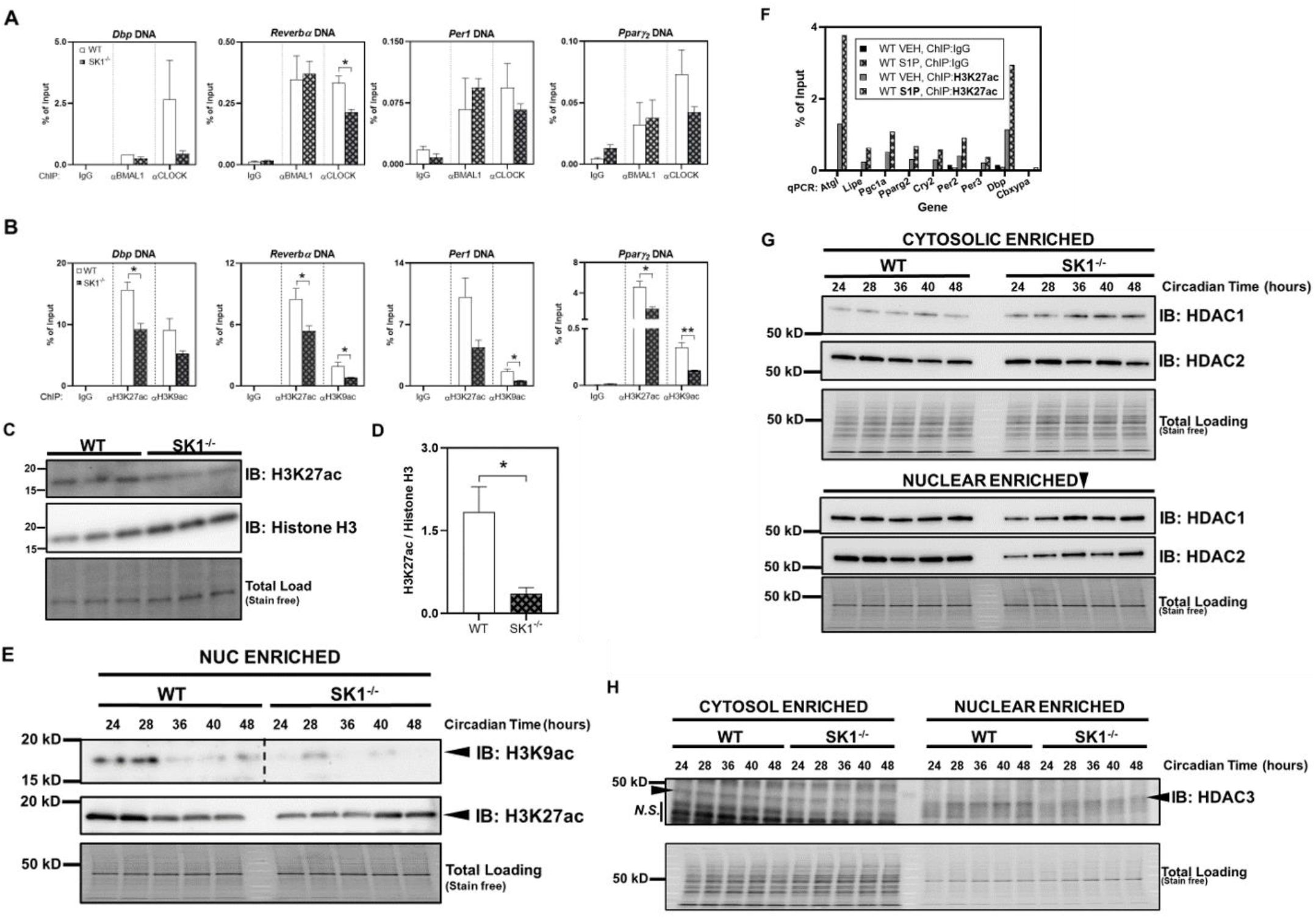
Loss of actively transcribing chromatin and CLOCK occupancy in SPHK1^-/-^ adipocytes and disruption of chromatin modifiers. **A**. WT and SPHK1^-/-^ primary adipocytes were synchronized and harvested at CT20 for ChIP of BMAL1 and CLOCK at the promoters of Per1 (negative core loop target), Reverbα (negative accessory loop target), Dbp (canonical/traditional, non-core-clock-mechanism BMAL1 target), and Pparγ (known circadian transcription factor, involved in anabolic adipocyte lipogenesis, BMAL1 target). (See Figure 2, page 13.) **B**. A mark of active transcription, acetylated histone 3 at lysines 9 and 27 (H3K9ac and H3K27ac) was immunoprecipitated at the DNA loci of Per1, Reverbα, Dbp, and Pparγ in synchronized WT and SPHK1^-/-^ primary matured adipocytes at CT20. **C**. Western blot analysis of H3K27ac in matured primary adipocytes from sWAT of WT and SPHK1^-/-^ mice. **D**. Quantification of the Western blot in (C.), displayed as relative acetylated histone3 K27 densitometry over total histone H3. **E**. Nuclear-enriched fraction of synchronized matured primary WT and SPHK1^-/-^ adipocytes showing acetylated H3K9 and H3K27 expression over 24 hours. **F**. WT synchronized primary matured adipocytes were treated with 150 nM S1P for 30 minutes and cells were subjected to ChIP of H3K27ac at the promoters for Atgl, Lipe, Pgc1α, Pparγ2, Cry2, Per2, Per3, Dbp, and at negative control Cbxypa (a marker present in pancreas tissue, but not adipose tissue). **G**. Cytosolic- and nuclear-enriched fractions of synchronized primary WT and SPHK1^-/-^ adipocytes and Western analysis of HDAC1 and HDAC2 proteins over 24 hours. **H**. Cytosolic- and nuclear-enriched fractions of synchronized primary WT and SPHK1^-/-^ adipocytes and Western analysis of HDAC3 over 24 hours. Experiments were performed 3 times in 3 separate preparations of primary cells for culture, and representative blots and ChIP batches are shown for the trends that were observed. *p < 0.05. Atgl, adipose triglyceride lipase; Cbxypa, carboxypeptidase A; Cry2, cryptochrome 2; Dbp, D-site of albumin promoter binding protein); Lipe, hormone sensitive lipase; Pgc1α, peroxisome proliferator-activated receptor gamma coactivator 1 alpha; Pparγ2, peroxisome proliferator-activated receptor gamma 2; Per2, period 2; Per3, period 3.

Histone markers associated with active chromatin (H3K9Ac, H3K27Ac) are known to exhibit circadian patterns, leading to assessment of these acetylated histone markers (Baerenfaller, et al., 2016; Cox and Takahashi, 2019; Feng, et al., 2011; Kim, et al., 2018; Sato, et al., 2017). H3K27Ac, served as a positive control when assessing the relatively low-expressed clock transcription factors, BMAL1 and CLOCK (Figure 5A). Intriguingly, both H3K9Ac and H3K27Ac occupancy signals at the *Dbp, Reverbα, Per1*, and *Pparγ*_*2*_ promoters were lower in SK1^-/-^ compared to control (Figure 5B). This prompted an examination of H3K27Ac protein expression at the same timepoint as the ChIP assay (CT20). H3K27Ac protein levels were markedly reduced in SphK1^-/-^ adipocytes relative to controls (Figure 5C-D), consistent with decreased acetylated histones at clock targets in SphK1^-/-^ cells. Further, assessment of acetylation patterns over a circadian time course revealed that H3K9Ac was nearly absent, while H3K27Ac expression was abundant in SphK1^-/-^ adipocytes (Figure 5E). This suggests absence of SphK1, typically located in the cytosol, has significant effects on nuclear transcriptional and epigenetic regulation.

Since S1P affects chromatin-associated histones in the nucleus, adipocytes were exogenously treated with S1P and ChIP of H3K27Ac was performed, followed by mRNA analysis of several clock and adipocyte targets. Lipolysis genes *Atgl* and *Lipe*, lipogenic genes *Pparγ*_*2*_ and *Pgc1α*, and circadian clock genes *Cry2, Dbp, Per2*, and *Per3*, acetylated histone H3 signals increased with S1P treatment compared to the albumin carrier alone (Figure 5F). These observations suggest that S1P facilitates a chromatin state that is favorable for transcriptional activation.

To further study the effects of S1P on chromatin modifiers, WT and SK1^-/-^ adipocytes were fractionated into cytosolic- and nuclear-enriched fractions. SK1^-/-^ adipocytes had increased cytosolic expression of HDAC1, paired with decreased nuclear HDAC1 compared to control (Figure 5G). Although cytosolic HDAC2 expression remained unchanged between genotypes, nuclear HDAC2 was reduced in SK1^-/-^ adipocytes compared to control, implicating SphK1 in the nuclear/cytoplasmic localization of Class I HDACs, potentially impacting their activity. Moreover, nuclear accumulation of HDAC3, was depleted in the nuclei of SK1^-/-^ adipocytes compared to control (Figure 5H). While decreased histone acetylation in SK1^-/-^ adipocytes suggest higher HDAC activity, this hypothesis remains to be tested, and no increased nuclear HDAC expression was found.

## Discussion

In conclusion, the work here demonstrated that genetic depletion of SphK1 led to defects in the adipocyte circadian clock with concomitant perturbations to the behavior of histones at key adipocyte and circadian genetic loci. Moreover, we propose that *Sphk1* is a circadian gene in adipocytes, due to the expression profile of *Sphk1* mRNA over a circadian time course in wildtype primary adipocytes, and the presence of clock transcription factors BMAL1:CLOCK at the promoter E-box sites in SphK1. Further investigation is needed to comprehensively decipher the role of SphK1/S1P as well as other sphingolipids in nuclear events such as homeostatic circadian transcription.

With respect to the cyclical nature of reversible or counterregulatory mechanisms, histone modifications (acetylation and deacetylation) and their modifiers are disrupted due to loss of SphK1. Consistent with the previously established notion that S1P is an inhibitor of HDAC, we found that treatment with S1P could increase the acetylated histone H3 signal at several key adipocyte functional and circadian regulatory genes. It was expected that an SphK1-null system would therefore be rampant with HDAC activity with subsequently transcriptionally inactive chromatin. This was demonstrated by decreased acetylated protein at two histone residues (K9 and K27) in SphK1^-/-^ cells and decreased mRNA expression from immunoprecipitated histones at important adipocyte circadian loci. Thus, SphK1^-/-^ cells had lower overall acetylated histone signal at several circadian target genes, which is in line with evidence that S1P, the product of SphK1, is a histone deacetylase inhibitor (15). Future studies delineating a role for SphK1 and S1P in chromatin modifications should include ChIP-Sequencing as a method for a more comprehensive analysis.

On the other hand, while nuclear and cytoplasmic localization of HDAC1, 2, and 3 was aberrant, it did not suggest, that HDAC1, 2, and 3-based deacetylation were responsible for the observations made by ChIP of decreased acetylated histone in SphK1^-/-^ adipocytes. CLOCK is a histone acetyltransferase (HAT), and because we observed increased nuclear levels of CLOCK in SphK1^-/-^ adipocytes compared to control, this potentially suggests a regulatory effect of SphK1 on HATs (17, 18). Despite this, CLOCK occupancy at several key promoters for which it is known to bind, decreased for several clock target genes involved in adipocyte homeostasis. Histone deacetylase and acetyltransferase enzymatic activity assays should be utilized to determine the global effect of SphK1 and S1P on their activity. Moreover, other HDACs, may be more active, which could explain the decreased acetylated histones found at the circadian and adipocyte gene promoters (19).

Literature suggest fatty acids oscillate in the bloodstream, and fatty acids have been shown in numerous contexts to induce the sphingolipid *de novo* biosynthetic pathway and some of its enzymatic genes, thus providing one mechanistic explanation for the observed daytime-dependent changes in sphingolipids (10, 18, 20-25). Sphingolipidomics data in human skeletal muscle biopsies obtained over a time course of 24 hours were highly suggestive of diurnal regulation of various sphingolipid species (18). The sphingolipid biosynthetic pathway represents a novel avenue for manipulation by therapeutics coupled with strictly scheduled dosing, or, stated differently, administration of sphingolipid-modifying drugs coupled with chronotherapy.

Figure 6 highlights a potential model by which SphK1 and its product S1P may influence the circadian clock. Histone modifications are known to be circadian and these modifications influence whether genes are enhanced or repressed (12, 25). *Sphk1* mRNA expression and the variations in the production of ^17^C-S1P demonstrate the cyclical nature of this lipid and its kinase. Thus, a model to encompass these aspects shows that S1P may impart a layer of control upon the circadian clockwork mechanism at the epigenetic level. Importantly, this study highlights for the first time a novel loop by which a cycling bioactive lipid exerts some control over the periodic oscillations of core clock genes and epigenetic machinery.

**Figure 6.**
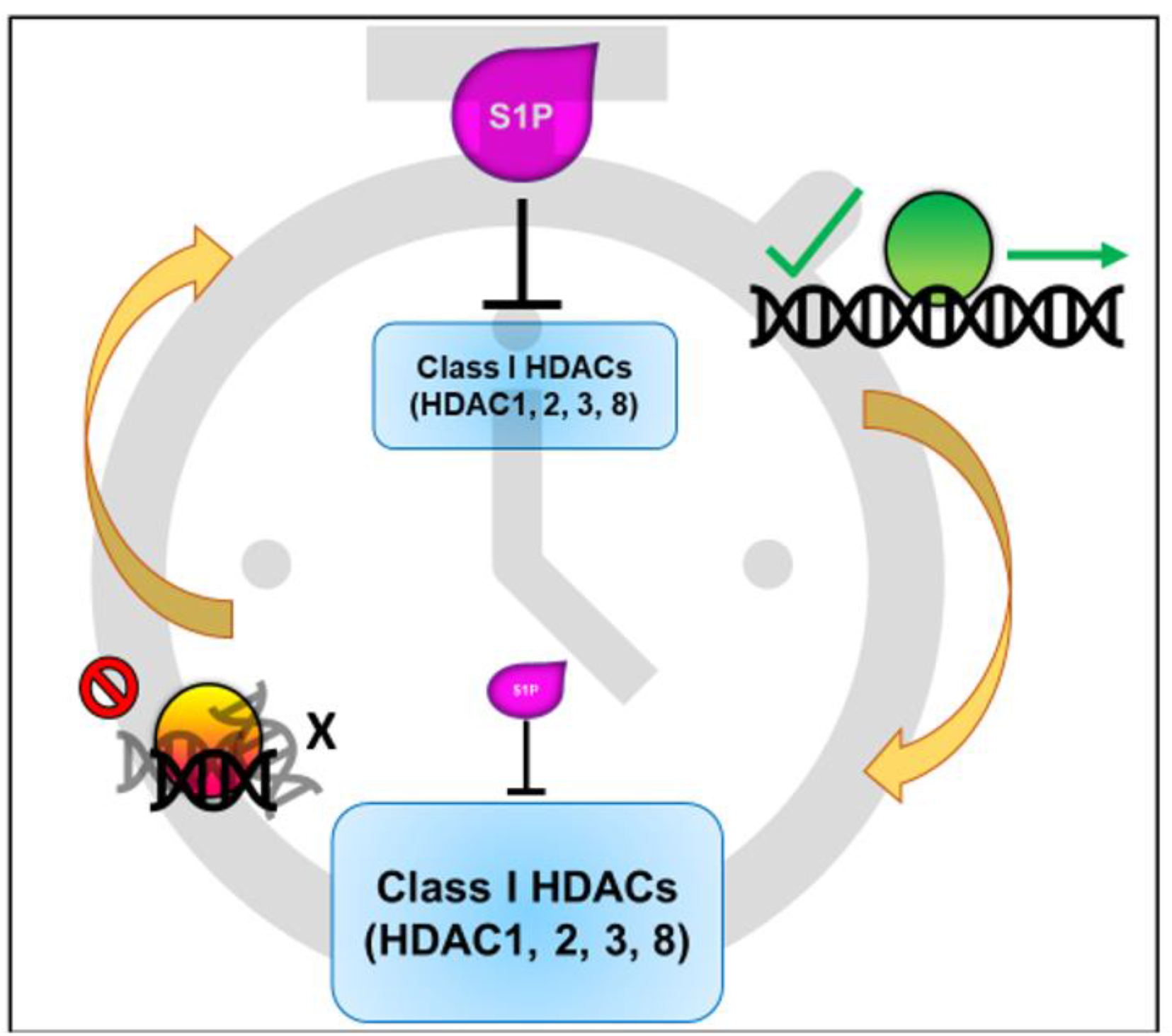
Proposed model of SPHK1/S1P regulation of adipocyte circadian rhythm. SPHK1 and S1P levels fluctuate over time, giving rise to a temporally-controlled inhibition on Class I HDAC activity. Temporal control of HDACs gives rise to chromatin conformations that enhance or repress transcription.

## Experimental Methods

### Cell Culture, Isolation of Stromal Vascular Fraction (SVF) and Differentiation to Mature Adipocytes

Primary adipose-derived stem cells (ADSCs) were isolated from inguinal adipose tissue of 3-8-week-old WT and SPHK1^-/-^ mice (Allende, et al., 2004). Subcutaneous adipose tissue was digested using 0.1% collagenase (Sigma, C6885) in digestion buffer (100 mM HEPES, 120 mM NaCl, 50 mM KCl, 5 mM glucose, 1 mM CaCl_2_, 1.5% BSA) in a 37° C shaking incubator (New Brunswick Scientific) (150 rpm) for 40 minutes. The digest was filtered through a 100-µM cell strainer and centrifuged at 500 ^×^ *g* for 5 minutes. The SVF pellet was re-suspended in media, filtered through a 40-µM cell strainer and plated. Media was changed after two hours to remove non-adherent cells. Cells were grown to confluency in proliferation media (DMEM/F12 50:50 (Corning, 10-090-CV), 10% fetal bovine serum (FBS) (Atlanta Biologicals), 1% penicillin streptomycin antimycotic cocktail (MilliporeSigma, A5955). 48 hours post-confluency, differentiation was induced using 10 µg/mL insulin (Santa Cruz Biotechnology, sc-360248), 1 µM dexamethasone (MilliporeSigma, D4902), 0.5 mM IBMX (Enzo Life Sciences, BMLPD1401000), and 1 µM rosiglitazone (Cayman Chemical, 71740) for two days.

Differentiation was maintained with insulin for days 3-4 and proliferation media days 5-8. Mature adipocytes were visualized under a brightfield microscope to assess lipid droplet formation, and plates with large lipid droplets and >90% differentiation were used for experiments (Figure 14).

### In vitro Circadian Rhythm

Unless otherwise noted, the *in vitro* circadian experiments are all performed in the same manner, in which primary WT and SPHK1^-/-^ ADSCs were brought to maturity, and on the day of the experiment, the cells were synchronized with 50% horse serum (Gibco, 26050-070) and 50% DMEM/F12 for 2 hours. Following the horse serum shock, the cell monolayer was gently rinsed once with warm 1x PBS and 2% fatty-acid-free BSA in DMEM/F12 was kept on the cells for the duration of the experiment, which generally lasted 48 hours *in toto*.

### S1P Treatment

WT primary matured, synchronized adipocytes were allowed to cycle for 24 hours, after which, cells were treated with a final concentration 150 nM S1P (Avanti Polar Lipids, 860492) complexed to 4 mg/mL FAF BSA or 4 mg/mL FAF BSA without S1P and assayed by ChIP for acetylated histone H3 signal at various adipocyte and circadian gene loci.

### ^17^C-Sphingosine Treatment

WT primary matured, synchronized adipocytes were allowed to cycle for 24 hours, after which, cells were treated with a final concentration 5 µM ^17^C-sphingosine (Avanti Polar Lipids, 860640) (dissolved in 100% ethanol) for exactly 30 minutes, collected immediately with addition of phosphatase inhibitors, for an exact volume of media in a 5-mL borosilicate glass tube and for a cell pellet in minimal PBS in a separate 5-mL borosilicate glass tube, capped, and stored at -80° C until analysis by tandem liquid chromatography and mass spectrometry lipidomics. This 30-minute procedure was repeated every 4 hours over a 24-hour period to assay *in situ* the enzymatic turnover of ^17^C-sphingosine to ^17^C-S1P (Spassieva, et al., 2007).

### Lipid Measurements

Sphingolipids were measured using liquid chromatography/tandem mass spectrometry at the Virginia Commonwealth University Lipidomics and Metabolomics Core and performed using previously published methods (Haynes, et al., 2009; Shaner, et al., 2009). ^17^C sphingolipids were measured at the Medical University of South Carolina Shared Lipidomics Resource according to previously published methods (Bielawski, et al., 2010; Spassieva, et al., 2007).

### Chromatin Immunoprecipitation (ChIP)

Primary WT and SPHK1^-/-^ matured, synchronized adipocytes were plated on 15-cm plates per 2 ChIPs. Briefly, cells were washed twice with room temperature 1x PBS and the dishes were then placed on an orbital rocker (Sigma, Z768502) in a measured volume of freshly-made PBS solution containing 2 mM disuccinimidyl glutarate (DSG) (ProteoChem, c1104) for 10 minutes. After 10 minutes, fresh 37% formaldehyde (Fisher, BP531-500) was added directly to the solution to a final concentration of 1% and continued swirling gently for another 12 minutes. Finally, the crosslinking reactions were quenched with glycine at a final concentration of 125 mM for 5 additional minutes. Following, the solution was discarded as hazardous waste and the cells were washed twice with cold 1x PBS. The PBS was aspirated, and the cells were scraped into 1.8 mLs adipocyte hypotonic lysis buffer (5mM PIPES pH 7.8, 10 mM KCl, 1% Igepal, plus protease/phosphatase inhibitors and PMSF immediately before use) per 15-cm plate. The cells were collected in a Dounce homogenizer and were sheared with 10 strokes of the Dounce until visibly dissociated. The cells were also passed through a 22G needle to ensure disruption of the monolayer but preservation of the nuclei. The resulting slurry was vortexed and kept on ice for 10 minutes. Next, the tubes were centrifuged at 8,500 ^×^ *g* at 4° C for 8 minutes to remove a significant fat cake at the top as well as the cytoplasmic infranatant and to leave the nuclear pellet intact and undisturbed. The pellets were resuspended in the same lysis buffer to wash excess adipocyte lipid from the nuclear pellet and vortexed. They were re spun, and the pellets were saved. Next, the pellets were suspended in 600 µL sonication lysis buffer (50 mM Tris-HCl, pH 8.0, 10 mM EDTA, 1% SDS, and protease/phosphatase inhibitors and PMSF fresh) per 15-cm plate. The 600 µL was split into two 1.5-mL Eppendorf tubes of 300 µL, and then were sonicated (Branson 150, Branson Ultrasonics, Danbury, CT) for 8 rounds of 30 seconds on, 30 seconds off in a 4° C water bath. The resulting sonicate was then subjected to centrifugation at 18,000 ^×^ *g* at 4° C to clarify the chromatin. The chromatin supernatant was collected and split to two chromatin immunoprecipitations as well as the 10% input control. Thus, approximately 500 µL of recovered sonicate was split to 200 µL per ChIP plus a 20-µL 10% input. The ChIP samples were diluted 1:10 with Dilution/Low Salt buffer (20 mM Tris-HCl pH 8.0, 150 mM NaCl, 2 mM EDTA, 1% Triton X-100, 0.1% SDS), pre-cleared with ChIP-grade Protein A/G magnetic beads (Thermo Fisher Scientific, 26162) for one hour, and immunoprecipitated with 1 µg of antibody overnight at 4° C while rocking. The following day, pre-equilibrated magnetic ChIP-grade Protein A/G beads were added to the samples for 2 hours. They were then subjected to extensive washing on the magnetic rack, performing two washes in Low Salt Buffer, 2 subsequent washes in high salt buffer (20 mM Tris-HCl pH 8.0, 500 mM NaCl, 2 mM EDTA, 1% Triton X-100, 0.1% SDS), 2 subsequent washes in LiCl Buffer (10 mM Tris-HCl pH 8.0, 500 mM NaCl, 1 mM EDTA, 1% deoxycholate, 1% NP-40, 0.25 M LiCl), and finally 2 washes in TE buffer (10 mM Tris-HCl pH 8.0, 1 mM EDTA). After these 8 washes, 100 µL of ChIP elution buffer (0.1 M NaHCO_3_, 1% SDS) was added to the beads, in addition to the reverse crosslinking reagent, 4 µL of 5 M NaCl, and these were incubated overnight at 65° C. The next morning, the samples were treated with RNase A (Thermo Fisher Scientific, EN0531) for 1 hour at 37° C, and after that, treated with Proteinase K (Thermo Fisher Scientific, EO0491) for 1 hour at 60° C. After this, the samples were subjected to DNA isolation and purification with a column-based washing approach (Zymo Research, D5205). The samples were eluted from the column in 120 µL and 2 µL per well, assayed in triplicate, were used for real time qPCR of the DNA.

**Table 1:**
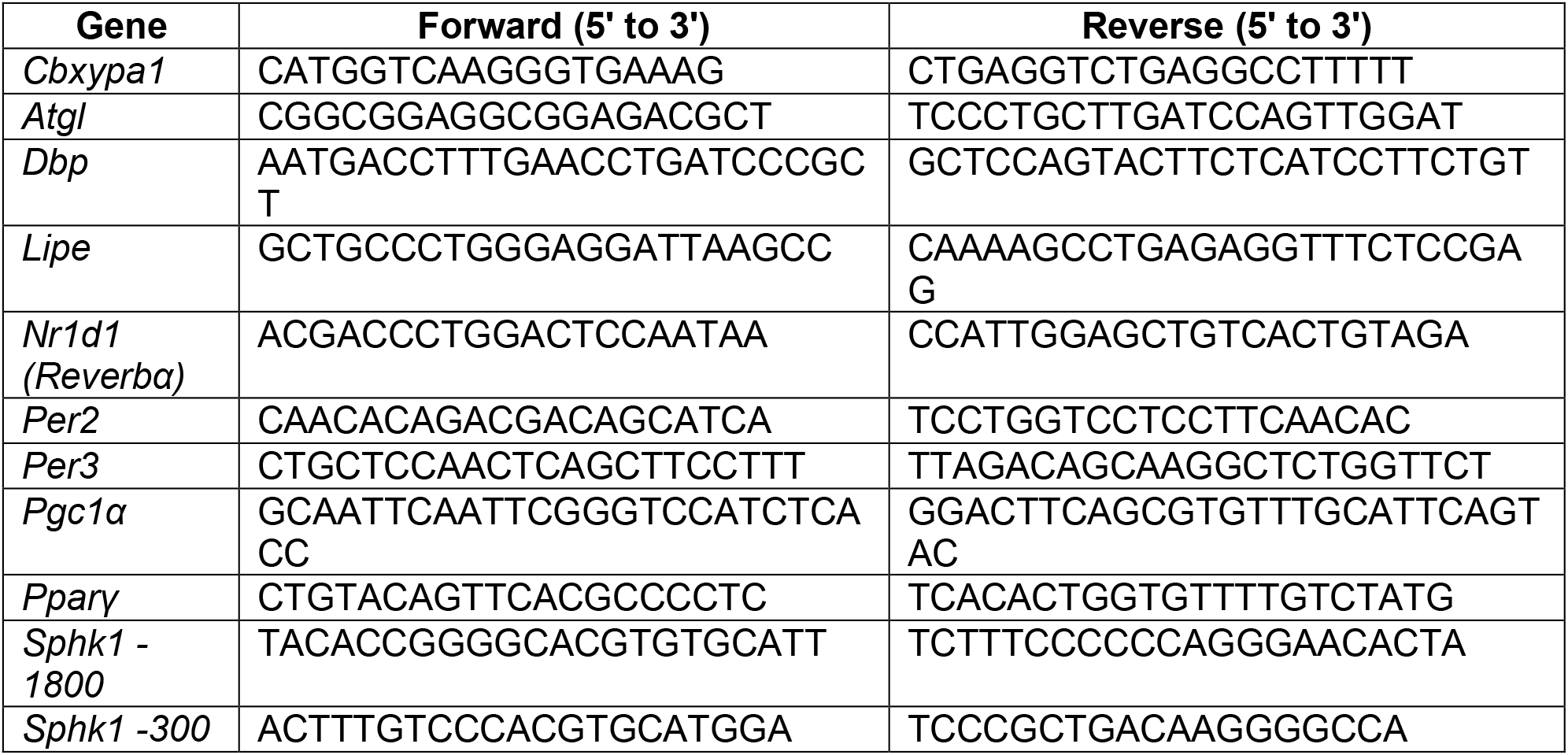
Primers used for ChIP are shown below:

**Table 2:** Antibodies used for ChIP are shown below:

| Application | Antibody | Catalog # | Concentration |
| --- | --- | --- | --- |
| ChIP | H3K9Ac | Thermo 710293 | 1 µg per IP |
| ChIP | H3K27Ac | Thermo MA5-24671 | 1 µg per IP |
| ChIP | IgG | Bethyl P120-101 | 1 µg per IP |
| ChIP and WB | BMAL1 | Bethyl A302-616A | 1 µg per IP |
| ChIP and WB | CLOCK | Bethyl A302-618A | 1 µg per IP |

### qPCR

Total RNA was isolated from cultured primary adipocytes or gonadal adipose tissue homogenized in Trizol (Invitrogen, 15596026) followed by RNeasy mini kit (Qiagen, 74106) extraction and column purification. cDNA was synthesized from 1 μg of total RNA using iScript Advanced cDNA Synthesis Kit (Bio-Rad, 1708890). Real time PCR was performed using a CFX96 Real-Time System (Bio-Rad) and SSoAdvanced Sybr (Bio-Rad, 172-5272). The primers used for cDNA qPCR and DNA CHIP-qPCR are shown in the following tables. Mean normalized expression was calculated by normalizing to the geometric mean of reference genes *Ppia* and *Tbp* in gonadal adipose tissue (*i*.*e*., root_2_[C_q_ gene 1 ⨯ C_q_ gene 2]) using the 2^-ΔΔCt^ method. ChIP normalization was carried out with a 10% input control. Primers for ChIP were tabulated in the *ChIP* section.

**Table 3:**
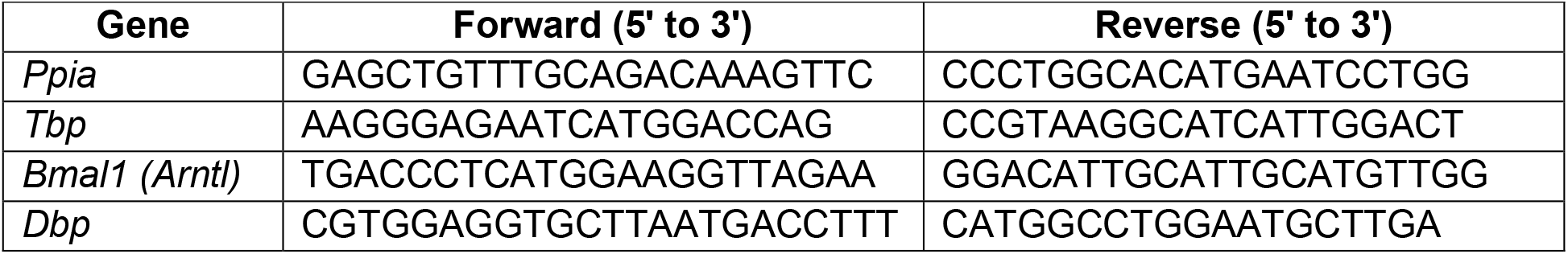
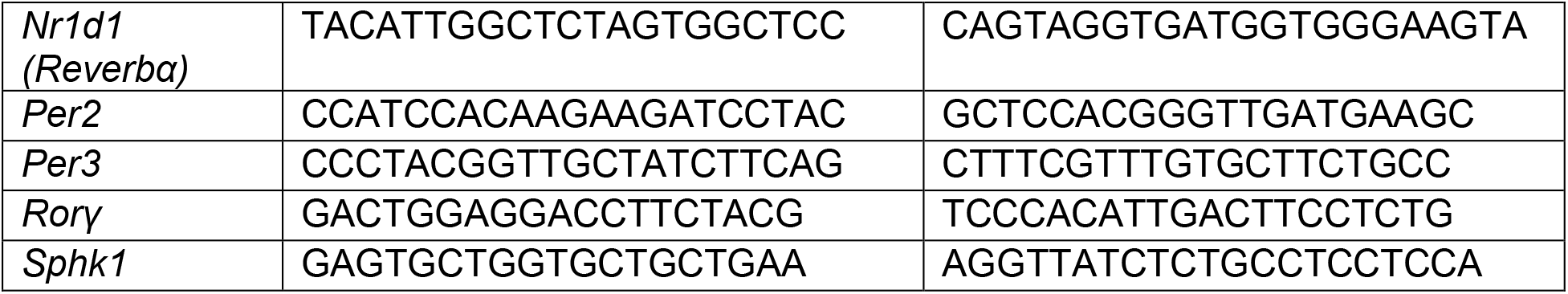
Primers used for qPCR are shown below.

### Cellular Fractionation for Nuclear and Cytoplasmic Enrichment

The Thermo NE-PER Kit (Thermo Fisher Scientific, 78835) was utilized to separate cytoplasmic (post-nuclear) and nuclear fractions from matured primary white adipocytes, with modifications. In a 10-cm dish, adherent mature adipocytes were gently rinsed twice with 1x PBS on the dish, and excess was aspirated with a 200-µL plastic pipet tip from the tilted dish. Next, ∼400 µL of CER I solution was added to the monolayer, and adipocytes were scraped into the solution. Importantly, the collected cells were then passed through a 22G syringe 5 times to disperse the cell conglomerate but keep the nuclei intact. These were vortexed and placed on ice for 10 minutes. Next, per the NE-PER protocol, 22 µL of CER II solution was added and tube was vortexed vigorously and placed on ice for 2 minutes. The tube was centrifuged at 500 × *g* for 5 minutes at 4° C. The fat cake and infranatant were collected to a separate tube (as the cytoplasmic/post-nuclear and fat cake components) while not disturbing the pellet at the bottom. The cytoplasmic tube was spun again in the same manner so that the fat cake and cytoplasmic-enriched infranatant could be separated. The cytoplasmic/post-nuclear infranatant was collected with a 28.5G insulin syringe, avoiding as much lipid as possible, at which point this sample was ready for BCA protein concentration determination. The tube with the pellet was cleaned of excess lipid on the sides of the tube by wicking carefully with a Kimwipe (Kimberly-Clark Professional, 34120), and the pellet was resuspended in 200 µL of NER, passed 5 times through a 28.5 gauge insulin syringe to break up the pellet, sonicated for 20 seconds in a sonicator bath, vortexed vigorously for 30 seconds, and placed on ice for 40 minutes. Finally, the nuclear natant was collected as the nuclear-enriched fraction after a 20-minute spin at 4° C at 18,000 × *g*. Cytoplasmic and nuclear fractions were assessed by Western blot after normalization by BCA. Nuclear fractions tended to be at least 5-fold lower in protein concentration.

### Western Blotting

Gonadal adipose tissue was homogenized in RIPA buffer with protease and phosphatase inhibitors (Thermo Fisher Scientific, 78446) using a Dounce homogenizer for at least 30 pulses, or until visibly homogenized. Homogenates were vortexed well and centrifuged at 10,000 ⨯ g for 10 minutes at 4° C. The resulting infranatant (below fat cake, and above cell debris pellet) was transferred to a new tube with an insulin syringe (28.5G). Cultured adipocyte whole-cell homogenates were homogenized in RIPA buffer with 1 mM PMSF, and protease and phosphatase inhibitors (Thermo Fisher Scientific, 78446). Cultured adipocyte cell fractions were collected as described in *Cellular Fractionation for Nuclear and Cytoplasmic Enrichment*. Protein content was quantified using a BCA protein determination assay (Thermo Fisher Scientific, 23225). 5-10 µg protein was used for western blotting. gWAT whole-cell homogenates, cultured adipocyte whole-cell homogenates, and fractionated adipocyte proteins were separated by SDS-PAGE, and transferred to PVDF membranes. The membranes were blocked for 1 hour in 5% BSA. Proteins were detected using HRP-linked anti rabbit secondary (1:5000; Cell Signaling Technology, 7074), Clarity ECL Western Blotting Substrate (Bio-Rad, 1705061) for HRP, and a ChemiDoc Imaging System (Bio-Rad, 17001401, 17001402). Blots were assessed for protein expression of proteins shown in the table on the following page. Vinculin (1:2,000; Cell Signaling Technology, 4650), Ponceau S, and stain-free total protein were used to determine even loading. Band intensity was quantified using ImageJ.

**Table 4:** Antibodies used for Western blotting analysis.

| Target | Antibody | Host | Supplier | Catalog # | Concentration or Dilution |
| --- | --- | --- | --- | --- | --- |
| Mouse | HDAC1 | Rabbit | Bethyl | A300-713A | 1:2,000 |
| Mouse | HDAC2 | Rabbit | Bethyl | A300-705A | 1:2,000 |
| Mouse | HDAC3 | Rabbit | Bethyl | A300-464A | 1:5,000 |
| Mouse | Histone H3 | Rabbit | Cell Signaling Technology | 4499 | 1:2,000 |
| Mouse | Lamin B1 | Rabbit | Cell Signaling Technology | 13435 | 1:1,000 |
| Mouse | Vinculin | Rabbit | Cell Signaling Technology | 4650 | 1:1,000 |
| Rabbit | HRP-anti-Rabbit IgG | Goat | Cell Signaling Technology | 7074 | 1:5,000 |
| Mouse | BMAL1 | Rabbit | Bethyl | A302-616A | 1 µg per IP |
| Mouse | CLOCK | Rabbit | Bethyl | A302-618A | 1 µg per IP |

### Mice

Adipocyte-specific *Sphk1* knockout (SK1^fatKO^) and WT C57BL/6 mice were utilized for *in vivo* studies (Anderson, et al., 2020). WT and SPHK1^-/-^ C57BL/6 mice were utilized for primary cell culture preparations for adipocytes (Allende, et al., 2004; Pappu, et al., 2007).

### Animal Model

All animal experiments conformed to the Guide for the Care and Use of Laboratory Animals and were in accordance with Public Health Service/National Institutes of Health guidelines for laboratory animal usage. The experimental groups consisted of male WT and SK1^fatKO^ C57BL/6 mice. Mice were housed in the animal facility at the Medical University of South Carolina. Standard chow and water were provided ad libitum, except when fasting was required (e.g., glucose tolerance testing). Animals were maintained on a 12h:12h light:dark cycle and ambient temperature was steadily 21° C. For diurnal experiments, unfasted WT and SK1^fatKO^ mice were sacrificed at ZT0, 3, 6, 9, 12, 15, 18, and 21, which corresponds to 6 a.m., 9 a.m., 12 p.m., 3 p.m., 6 p.m., 9 p.m., 12 a.m., and 3 a.m. At the time of sacrifice, mice were humanely euthanized by isoflurane (Hospira, Inc., Lake Forest, IL) followed by cardiac puncture. Cardiac blood was prepared for non-hemolyzed serum, aliquoted, and stored at -80° C until further analysis. Tissues were collected accordingly as fresh snap frozen in liquid nitrogen and stored at -80° C until further processing. All experiments were performed under clean conditions, were approved by the Medical University of South Carolina Institutional Animal Care and Use Committee, the Virginia Commonwealth University Institutional Animal Care and Use Committee, the Ralph H. Johnson Veterans Affairs Medical Center, and the Hunter Holmes McGuire Veterans Affairs Medical Center.

### Statistical Analysis

All values are presented as mean ± SEM. For single pairwise comparisons of normally distributed data sets, a Student’s *t*-test was performed. For multiple comparisons of means, a one-way ANOVA with Tukey-Kramer post hoc test was performed. *p*<0.05 was considered statistically significant. All hypothesis tests were conducted using Graphpad Prism 8 software.

## Acknowledgements

This study and its personnel were supported in part by grants to Dr. Cowart from the National Institutes of Health (NIH; R01HL117233 and R01HL151243) and Veterans’ Affairs (IKBX006315 and 5I01BX000200). Grants F31HL156529 and T32HL149645 were provided to Dr. Kovilakath; R21AA030647 to Drs. Cowart and Montefusco. Services and products in support of the research project were generated by the following Virginia Commonwealth University/Massey Comprehensive Cancer Center Shared Resources: The Lipidomics and Metabolomics Shared Resource, the Cancer Mouse Models Core, the Microscopy Shared Resource, and the Transgenic/Knockout Mouse Shared Resource, supported in part with funding from NIH-NCI Cancer Center Support grant P30 CA016059.

